# Aging, mortality and the fast growth trade-off of *Schizosaccharomyces pombe*

**DOI:** 10.1101/128298

**Authors:** Hidenori Nakaoka, Yuichi Wakamoto

**Author notes:** CONTACT (Y.W.).

## Abstract

Replicative aging has been demonstrated in asymmetrically dividing unicellular organisms, seemingly caused by unequal damage partitioning. Although asymmetric segregation and inheritance of potential aging factors also occurs in symmetrically dividing species, it nevertheless remains controversial whether this results in aging. Based on large-scale single-cell lineage data obtained by time-lapse microscopy with a microfluidic device, in this report, we demonstrate the absence of replicative aging in old-pole cell lineages of *Schizosaccharomyces pombe* cultured under constant favorable conditions. By monitoring more than 1,500 cell lineages in seven different culture conditions, we showed that both cell division and death rates are remarkably constant for at least 50–80 generations. Our measurements revealed that the death rate per cellular generation increases with division rate, pointing to a physiological trade-off with fast growth under balanced growth conditions. We also observed the formation and inheritance of Hsp104-associated protein aggregates, which are a potential aging factor in old-pole cell lineages, and found that these aggregates exhibited a tendency to preferentially remain at the old-poles for several generations. However, the aggregates were eventually segregated from old-pole cells upon cell division and probabilistically allocated to new-pole cells. The quantity and inheritance of protein aggregates increased neither cellular generation time nor cell death initiation rates. Furthermore, our results revealed that unusually large amounts of protein aggregates induced by oxidative stress exposure did not result in aging; old-pole cells resumed normal growth upon stress removal, despite the fact that most of them inherited significant quantities of aggregates. These results collectively indicate that protein aggregates are not a major determinant of cell fate in *S. pombe*, and thus cannot be an appropriate molecular marker or index for replicative aging under both favorable and stressful environmental conditions.

## Introduction

Replicative aging in unicellular organisms is defined by a gradual increase in generation time and probability of death as cell divisions increase. In cases of asymmetrically dividing unicellular organisms such as *Caulobacter crescentus, Saccharomyces cerevisiae*, and *Candida albicans*, aging is manifested and linked to morphological asymmetry [1–3]. The situation, however, is less clear for symmetrically dividing organisms. While some evidence suggests replicative aging in old-pole cell lineages of *Escherichia coli* [4–6], Wang *et al*. reported that growth rates of *E. coli* old-pole cells did not significantly alter over 200 generations, despite the gradual increases in filamentation and death rates [7]. For the symmetrically dividing fission yeast *Schizosaccharomyces pombe*, earlier studies suggested replicative aging by observation of asymmetry in cell volume at divisions followed by the deaths of the larger cells, and asymmetric segregation of carbonylated proteins (one of the biomarkers of oxidative stress). Additionally, it was suggested that inheritance of carbonylated proteins and a birth scar might inversely correlate with survival probability [8–10]. In a more recent study, however, Coelho *et al*. showed that potential aging factors such as an old-pole, a new spindle pole body, and protein aggregates did not correlate with generation time, suggesting that *S. pombe* does not age, at least under favorable conditions [11].

The key mechanism to generate aging lineages (and their rejuvenated counterparts) is thought to be asymmetric segregation of “aging factors”, regardless of the mode of cell division. Among the potential aging factors are aggregates of misfolded proteins [12,13]. In *E. coli*, naturally-occurring protein aggregates reside exclusively at old-pole ends, probably because of nucleoid occlusion, and a negative correlation between the aggregate burden and growth rate is observed [5,6]. Likewise, in *S. cerevisiae*, protein aggregates are preferentially found in aging mother cells. Active mechanisms that have been suggested to contribute to asymmetric segregation include organelle-associated confinement and actin cable-dependent retrograde flow [14–17]. Asymmetric damage segregation and its association with aging were also suggested in *S. pombe* cultured under heat or oxidative stress conditions [11]. Although it was previously shown how aggregates are formed, move, and segregate during cell division, it is not clearly established whether they contribute to increased death rate [11,18].

To study the aging process of yeasts, lineage tracking on an agar plate is conventionally performed [19]. This requires micromanipulation to remove daughter cells and is relatively labor intensive, thus precluding high-throughput and long-term analyses. An increasing number of studies at the single-cell level for various model organisms utilize microfluidic devices made of polydimethylsiloxane (PDMS), a chemically-stable and biocompatible silicone, in combination with automated microscopic imaging techniques [20–25]. One such device, termed the “Mother Machine”, was originally developed to track *E. coli* old-pole cell lineages with considerably higher throughput (10^5^ individual old-pole cells) and longer duration (200 generations) than previous studies [7]. Mother Machine-like microfluidic devices for *S. pombe* have been reported recently, and they demonstrated the absence of replicative aging in rich medium [26,27].

In this work, we measured more than 1,500 fission yeast old-pole cell lineages up to 80 generations using a custom-built Mother Machine-like microfluidic device. By measuring cell division and death rates in seven different balanced growth conditions, we confirmed the absence of replicative aging in old-pole cell lineages in all of the tested environments and found a positive correlation between the two rates. We observed formation, growth, inheritance, and asymmetric segregation of Hsp104-associated protein aggregate in the old-pole lineages and demonstrated that inheritance and quantity of protein aggregate affected neither generation time nor triggering of cell death. In addition, a large amount of protein aggregate induced by transient stress treatment could also be tolerated without affecting cellular growth rates. Collectively, our results suggest that protein aggregate does not serve as an aging marker under both favorable and stressful conditions.

## Results

### Old-pole cell lineages exhibit aging-free growth over tens of generations in a microfluidic device

We designed and developed a microfluidic device for long-term tracking of old-pole cell lineages of *S. pombe* (Fig. 1A, 1B, and Fig. S1). Our device has essentially the same architecture as the “Mother Machine”, which was originally developed by Wang *et al*. for studying aging and growth in *E. coli* [7], except that the dimensions of the internal channels were scaled-up for fission yeast, which are physically larger. During time-lapse experiments, the device was constantly supplied with fresh medium to keep the environmental conditions around the cells unchanged. We experimentally confirmed that the medium reached the ends of the observation channels within 5 min, both in the absence and presence of cells (Fig. S2 and Movie S1). Cells grew and divided aligned in the observation channels, and cells that spilled out from the observation channels to the trench were washed out by the flow of medium (Fig. 1B and Movie S2). These settings allowed us to follow the division dynamics of cells located at the ends of the observation channels, referred to as old-pole cells (or mother cells), typically for 50–80 generations. Time-series data on cell size (determined by visualized cell area) for every cell lineage were extracted from a set of time-lapse images (Fig. 1C). We analyzed more than 1500 single-cell lineages in each experiment, which is comparable to, or larger than, the numbers of cell lineages analyzed in similar microfluidic experiments [7,27].

**Fig 1.**
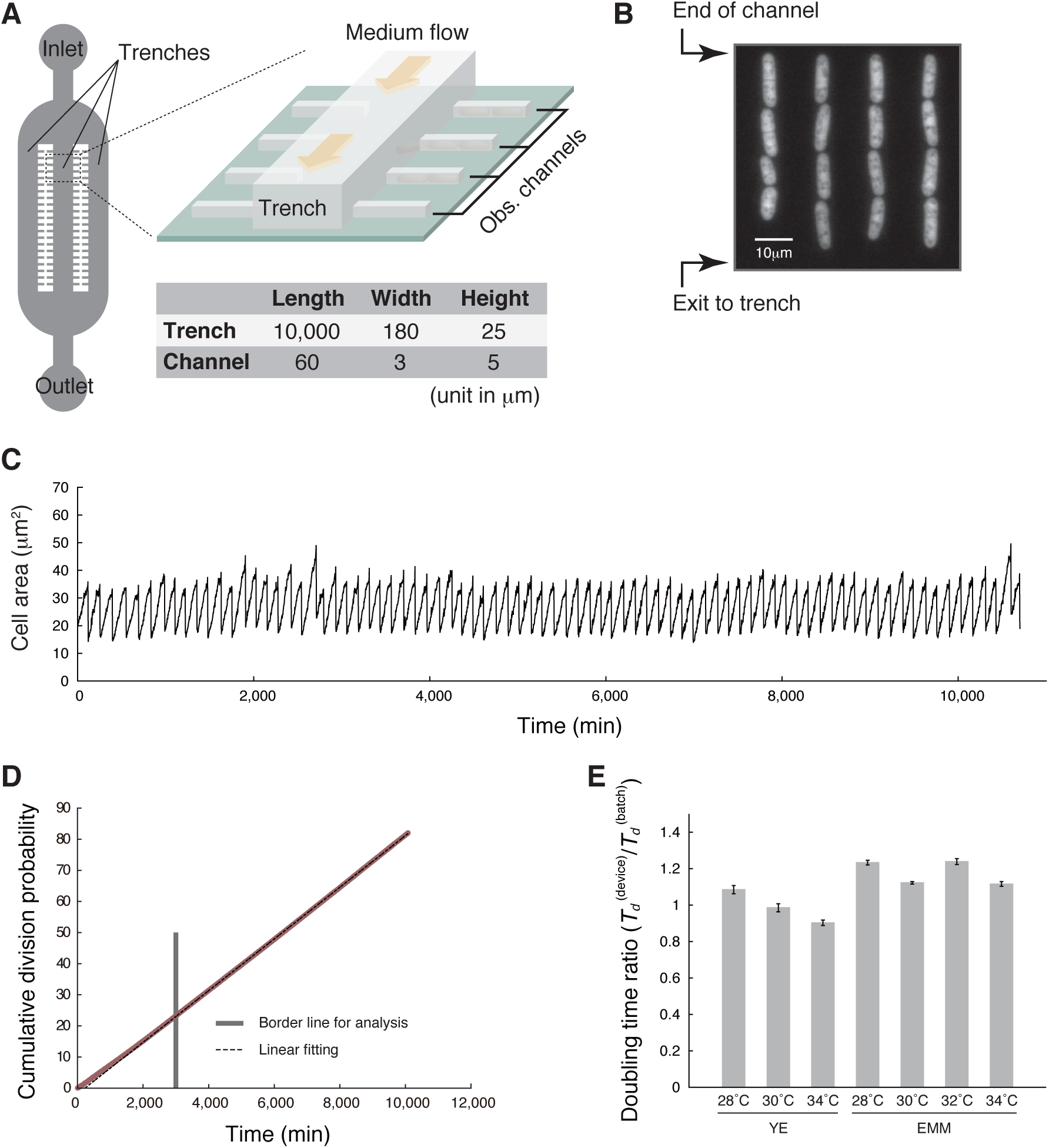
Stability of cell division rate in the microfluidic device. (A) Schematic representation of the device (not to scale, for clarity). Approximate dimensions of the trenches and observation channels are presented in the table. (B) An example of a fluorescence image of yeast cells expressing mVenus loaded into the observation channels. (C) Trajectory of cell size of a single representative lineage in YE at 34°C. (D) The cumulative division probability in YE at 34°C, plotted against time (red). Linear fitting (black broken line) was performed using the time window (*t* ≥ 3,000 min in this example, indicated by a gray vertical line) in which stable growth was achieved. See also Fig S3. (E) Estimated population doubling times in the microfluidic device (*T_d_* ^(device)^) relative to those in batch cultures (*T_d_* ^(batch)^). See Materials and Methods for estimation of the population doubling times from the generation time distributions.

We performed time-lapse experiments employing seven different culture conditions with different media (yeast extract medium [YE] or Edinburgh minimal medium [EMM]) and temperatures (see Table S1 for the summary of all measurements). Plotting the cumulative division probability against time confirmed that division rates were strikingly stable except during initial measuring (Fig. 1D and Fig. S3). This early instability in division rates reflected a lag in cell recovery from the slow-growing state that follows the loading of the cells into the microfluidic device (see Materials and Methods). Population doubling times calculated from the distributions of generation times (see [28–30] for reference) were close to those determined in batch culture experiments in the same media and temperature conditions (Fig 1E). This indicates that the medium exchange rate in the device is sufficiently high. The stability in division rates for 50–80 generations, in turn, suggests an absence of deterioration in the reproductive ability of old-pole cells, under favorable culture conditions.

Despite these favorable growth conditions, we observed the deaths of individual cells at low frequencies over the entire observation period (Fig. 2A, Movie S2, and Table S1). Because the death events were observed throughout the time-lapse experiments and in every imaged position, they were not caused by temporal and/or local alterations in culture environments. The behaviors of cells destined for death were heterogeneous, but could be broadly categorized into three types: Type I (swollen), Type II (hyper-elongated), or Type III (shrunken). Approximately 80% of the death events were categorized as Type I, and in almost all of these cases, siblings in the same observation channel synchronously died (Fig. S4D, E, and Movie S2). These observations are consistent with a recent report using a similar microfluidics system [27]. The synchronous deaths were also observed in another PDMS microfluidic device, where the observation channels accommodate greater numbers of cells than the Mother Machine. Importantly, we observed the synchronous deaths even when the dying siblings were spatially separated, whereas the other surrounding cells continued dividing normally (Fig. S4F and Movie S3). These findings suggest that the synchronous deaths are not induced by local environmental changes in channels, but triggered in their common ancestor cells.

**Fig 2.**
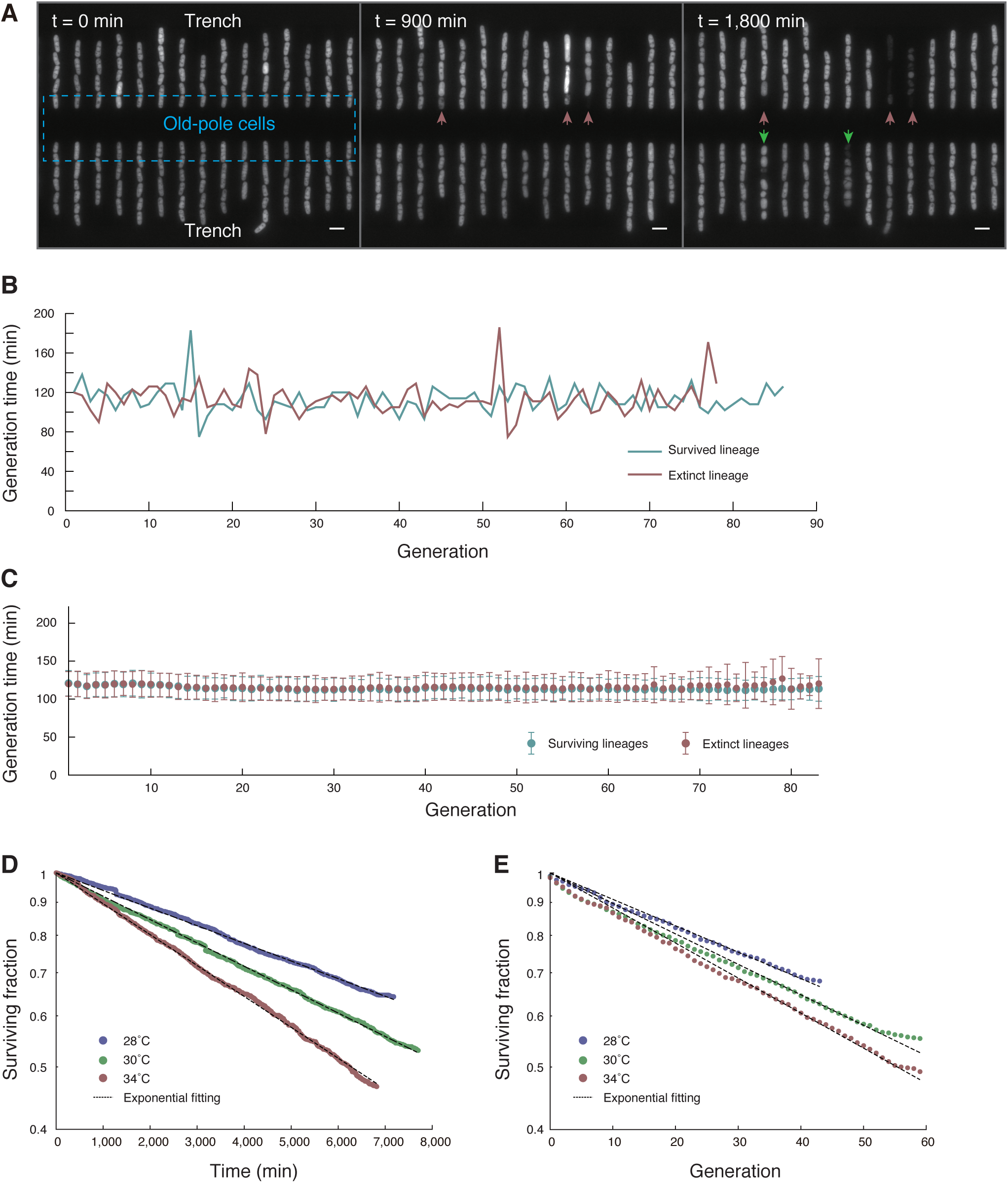
Characterization of cell deaths in constant environments. (A) Three fluorescence images of the same position at different times in YE at 34°C. While all of the old-pole cells were alive at the beginning (*t* = 0), three died by 900 min (red arrows), and two others by 1,800 min (green arrows). Scale bars indicate 10 μm. (B) Representative generation time transitions in surviving (blue) and extinct (red) lineages in YE at 34°C. (C) Transition of mean generation time for each generation in YE at 34°C. Error bars represent standard deviations. See also Fig S5. (D) Decay of the surviving fraction of old-pole cell lineages against time in YE at three temperature conditions (blue, 28°C; green, 30°C; and red, 34°C). See also Fig. S6. (E) Decay of surviving fractions plotted against generation count.

We did not detect any preceding progressive signatures in the growth and division histories of dead cells. For example, the transitions in generation times of the extinct cell lineages were indistinguishable from those of the surviving lineages; no obvious or discernible increase in generation times was observed prior to cell deaths (Fig. 2B). The stability of generation times in the extinct cell lineages was further confirmed by comparing the means and standard deviations of generation times for each generation between the surviving and extinct lineages (Fig. 2C, and Fig. S5). In addition, the number of surviving cell lineages decayed exponentially with time and generation count (Fig. 2D, 2E, and Fig. S6), indicating that cell deaths occurred randomly with fixed probabilities and that every lineage exhibited an equal chance of abrupt death. The death rates estimated from the decay curves were small, in the order of 10^-5^ per minute (or 10^-2^ per generation). We noticed that our standard fluorescence imaging conditions induced weak photo-damage [31]. Consequently, the estimated death rates were slightly higher than of those obtained by bright field imaging alone, but the constancy of the death rates was unaltered (Fig. S4B and S4C).

### Trade-off between reproduction and survival in balanced growth conditions

We next investigated how cellular division and death rates might be interrelated. As presented in Fig. 3A, we found that the death rate increased linearly with the division rate. The values of each data point (including error estimates) and the corresponding culture conditions are summarized in Table S3. For example, in YE at 34°C, the division rate was *r* = (8.94 ± 0.01) × 10^-3^ min^-1^ (mean generation time *τ*_b_ = 1/*r* = 112 min) and the death rate was *k* = (1.1 ± 0.1) × 10^-4^ min^-1^ (characteristic lifetime *τ*_d_ = 1/*k* = 9.1 × 10^3^ min = 6.3 day). Additionally, in EMM at 28°C, the division rate was *r* = (4.29 ± 0.01) × 10^-3^ min^-1^ (*τ*_b_ = 233 min) and the death rate was *k* = (2.0 ± 0.4) × 10^-5^ min^-1^ (*τ*_d_ = 5.0 × 10^4^ min = 35 days). Thus, fast growth significantly shortens the lifetime of single cells. Reformatting the plot reveals that the expected life span of single-cell lineages in units of generation (*τ*_d_/*τ*_b_) also decreases with division rate, asymptotically approaching the minimum bound (the shortest expected life span) of approximately 50 generations (Fig. 3B, gray broken line). The decrease in expected life span can be attributed to the fact that the death rate reaches zero with a positive division rate value (*r*^min^ = [3.6 ± 0.2] x 10^-3^ min^-1^, equivalently *τ*_b_^max^ = 280 min) as shown in Fig. 3A.

**Fig 3.**
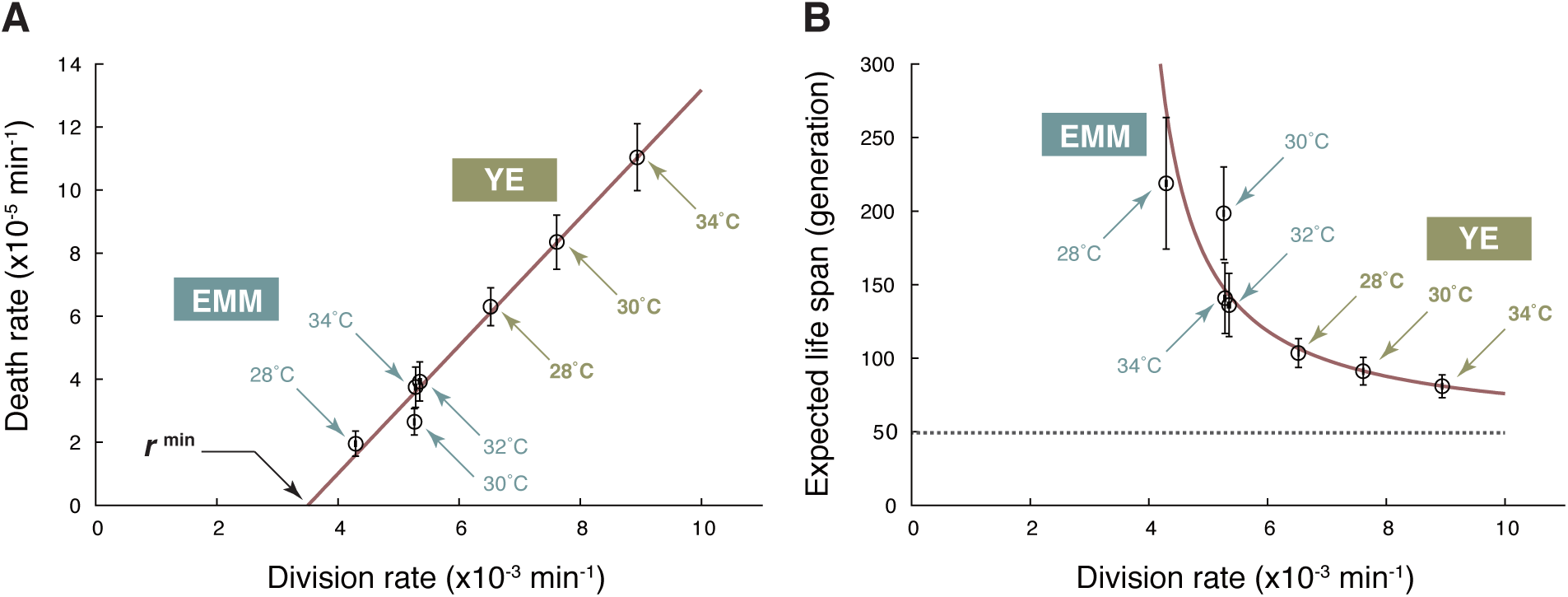
Trade-off between division and death rates. (A) Relationship between division rates (*r*) and death rates (*k*). The red line represents the best linear fit to the data points (open circles) by the least squares method. The error bars represent ±2 standard error ranges (see Materials and Methods for the rigorous definitions). (B) Relationship between division rate (*r*) and expected life span (τ_d_/τ_b_ = *r/k*) of a single-cell lineage. The red line is a theoretical curve based on linear fitting in (A): *k* = α (*r – r*^min^), where the slope α= (2.0 ± 0.1) × 10^-2^, and *r* ^min^ = (3.5 ± 0.2) × 10^-3^ min^-1^. The gray broken line represents the minimum expected life span (1/ α) in the fast-growth limit.

### Aggregated protein deposits remain in old-pole cell lineages for multiple generations but are eventually segregated to new-pole cells

Our observation that fission yeast old-pole cell lineages are unlikely to undergo replicative senescence motivated us to proceed to monitor long-term dynamics of protein aggregation to gain insight into how these lineages avoid aging. In general, protein aggregates are associated with molecular chaperones and heat shock proteins that can extricate protein monomers from the aggregates and refold them into their native structures [12,13]. One of the most well-studied heat shock proteins in yeasts is Hsp104, an ATP-dependent disaggregase that is often used as a molecular marker of protein aggregation in both *S. cerevisiae* and *S. pombe*. A strain that expresses Hsp104-GFP from the native chromosomal locus was observed in the microfluidic device for approximately 50 generations (Fig. 4A and Movie S4). The time-lapse imaging revealed that most healthy growing cells had zero or one major GFP focus. We defined aggregates as a set of connected pixels whose fluorescence (i.e. GFP) intensity exceeded a defined threshold value, and quantified aggregate amounts by integrating fluorescent intensity within the connected area (Fig. S7). The distribution of aggregate amounts was roughly exponential (Fig. 4B), which is consistent with previously reported results [18]. Fig. 4C presents a representative dynamic of formation, growth, and segregation of protein aggregates in an old-pole cell. Once formed at an old-pole end, the (major) aggregate grew and tended to remain at the pole for many generations, but it occasionally migrated toward the new-pole end, and was subsequently segregated to the new-pole cell (Movie S4), which is qualitatively consistent with an earlier report [18]. The distribution of aggregate inheritance duration, which is defined as the time interval between two successive “born-clean events” in units of generation, had a peak at four generations with an extended tail to the right and spreading over more than 40 generations (Fig. 4D). The tail can be approximately fitted by an exponential curve with a decay rate of λ = 0.13 (generation^-1^), suggesting that the segregation of protein aggregate to a new-pole cell is a random process that occurs once in every 1/λ = 7.8 generations on average. These results revealed that fission yeast old-pole cell lineages could escape from the burden of protein aggregate.

**Fig 4.**
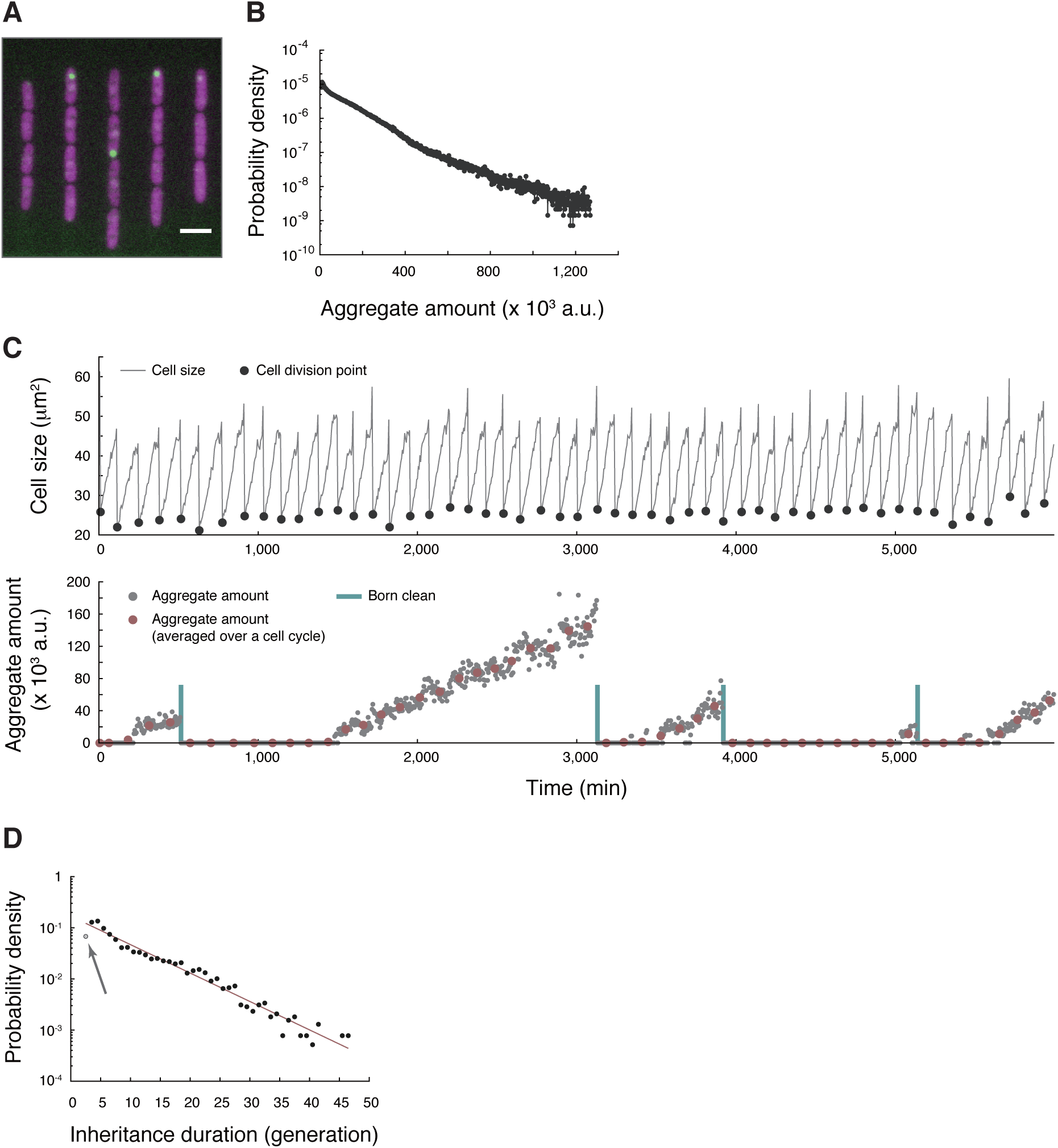
Formation and segregation dynamics of Hsp104-associated protein aggregate. (A) An example of merged fluorescence images (GFP and RFP channels) of the strain HN0045 cultured in the microfluidic device. Green: Hsp104-GFP. Magenta: mCherry. Scale bar indicates 10 μm. (B) Distribution of aggregate amounts; *N* = 932,801. (C) Typical cell size trajectory of HN0045 grown in the microfluidic device (top) and dynamics of formation, growth, and segregation of Hsp104-GFP foci in the lineage (bottom); gray closed circles: aggregate amount at each time point; red closed circles: cell-cycle-averaged aggregate amount; blue vertical lines: points of cell divisions that produced aggregate-free old-pole cells. A time interval between two adjacent blue lines is defined as duration of aggregate inheritance. (D) Distribution of aggregate inheritance interval. Data points, excluding the gray one indicated by an arrow, were fitted by shifted-exponential distribution: *p*(*x*) = *λe*^*−λ*(*x–μ*)^. A line of best fit (1/λ = 7.8 and μ = 2.1) is shown colored in red.

### Protein aggregation affects neither generation time nor initiation of cell death

Although the mean generation time of the old-pole cell lineages was stable (Fig. 2B and 2C), heterogeneity in each cell cycle length might be related to protein aggregation. We quantified the load of protein aggregation using two metrics: 1) aggregate amount; and 2) aggregate age, the latter being defined as elapsed time (in units of generation) since the last birth without aggregate inheritance (indicated by “Born clean” bars in Fig. 4C). The former evaluates the current load of aggregation, whereas the latter evaluates the burden of possessing the aggregate for prolonged periods. We first simply plotted generation time against (cell cycle-averaged) aggregate amount (Fig. 5A) and aggregation age (Fig. 5B), detecting no correlations.

**Fig 5.**
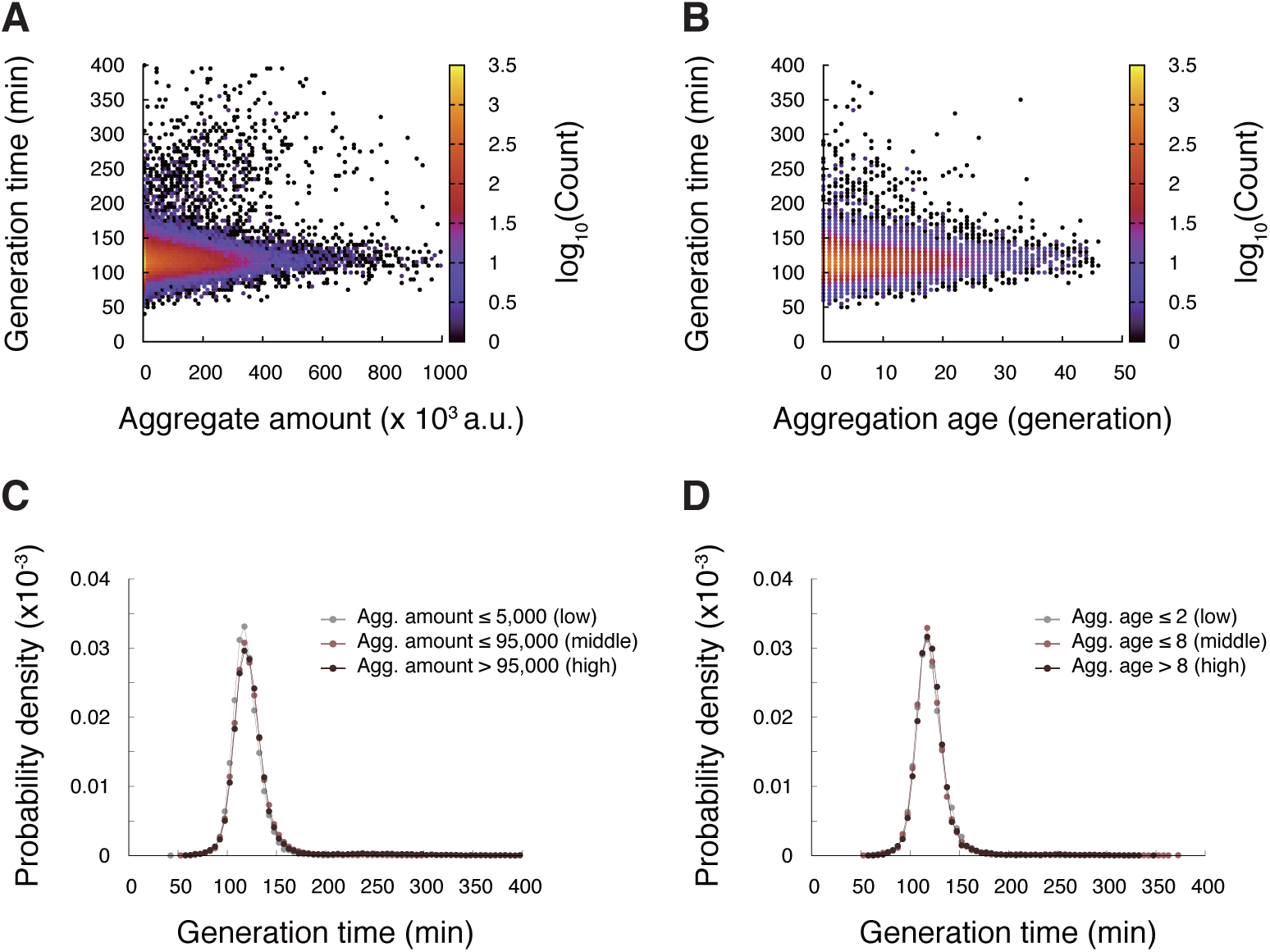
Hsp104-associated protein aggregates do not affect generation time. (A) Correlation between aggregation level and generation time; *N* = 53,683 and Spearman’s ρ = 0.081. (B) Correlation between aggregation age and generation time. Data count associated with each point is indicated by colors; *N* = 39,898 and Spearman’s rank-order correlation coefficient ρ = 0.022. (C) Generation time distributions for different levels of aggregate amount. *N* = 15,551 (aggregate amount ≤ 5,000), 20,438 (5,000 < aggregate amount ≤ 95,000), and 17,806 (aggregate amount > 95,000), respectively. (D) Generation time distributions for different levels of aggregation age. *N* = 13,481 (aggregate age ≤ 2), 14,481 (2 < aggregate age ≤ 8), and 13,115 (aggregate age > 8), respectively.

To analyze such relations in greater detail, we partitioned the data points in Fig. 5A and 5B into three classes (low, middle, and high) according to the aggregation metrics (aggregate amount or aggregation age), and compared the generation time distributions among the classes (Fig. 5C and 5D). The distributions were essentially identical among the classes for both aggregation indices, which strongly indicates that cell cycle length is unaffected by protein aggregation.

Next, we examined if the amount and inheritance of protein aggregation trigger cell death. Fig. 6A illustrates protein aggregation dynamics (Hsp104-GFP aggregate amount) along with the level of constitutively expressed protein (mCherry mean fluorescence intensity corresponding to its cellular concentration) for both survived and extinct lineages. Typically, cells destined for death exhibited accelerated accumulation of protein aggregates immediately prior to death (Fig. 6A [top panel], after 4,500 min); we detected accelerated accumulation in 79% (427 out of 541) of extinct lineages. The commencement of accelerated accumulation was detectable by clear kinks in the transitions of aggregate amounts. These kinks seem to identify the time of initiation of cell death because other apparent functional deteriorations, such as radical increases in mCherry expression and changes of cellular morphology, also initiated concurrently (Fig. 6A, 6B, and Movie S4). Interestingly, even after the onset of these abnormalities, cell division occurred a few times before death (Fig. 6C), which might underlie the observed synchronized deaths (Fig. S4D, S4F, and S4G).

**Fig 6.**
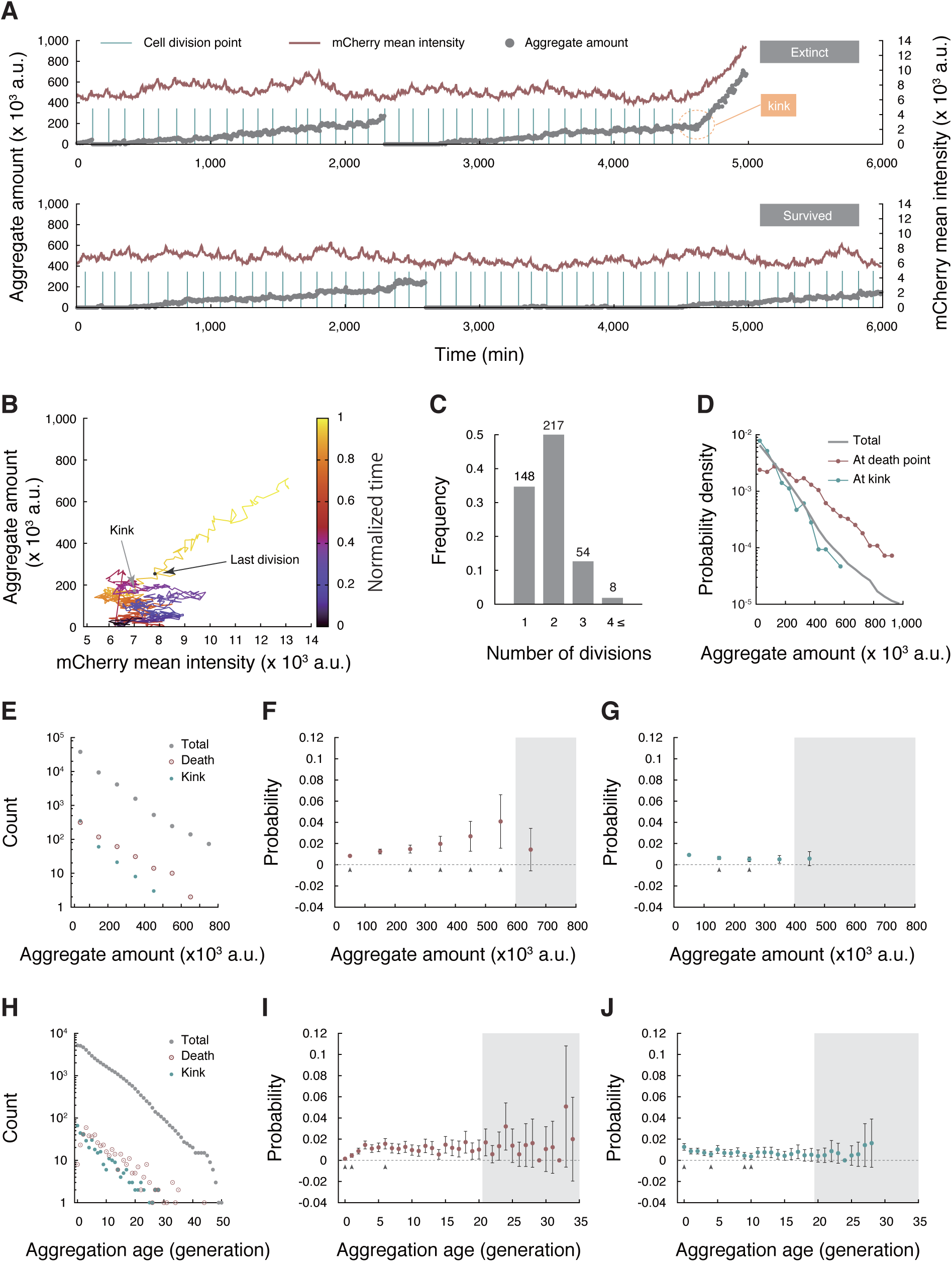
Relationship between Hsp104-associated protein aggregate and death. (A) Typical dynamics of Hsp104-associated protein aggregate (gray closed circles) and constitutively expressed mCherry level (red line) of an extinct lineage (top) and a survived lineage (bottom). The dying process begins around 4,500 min (indicated by a dotted orange circle) in this example, after which both aggregate amount and mCherry signals increase. (B) Dynamics of protein aggregation and mCherry levels plotted in a 2D plane. The data are from the same (extinct) cell lineage shown in (A). Time evolution is indicated by colors from dark blue to yellow. The black and gray points indicate the states at the last division and the kink position, respectively. (C) The number of cell divisions after the kinks until cell death. Figures on top of the bars represent the number of lineages identified. (D) Distributions of aggregate amount at death points (red), kinks (blue), and for all ROIs [Region Of Interest] (gray). (E) Distributions of aggregate amount at birth for all identified cell cycles (gray), for the last generations in extinct lineages (red), and for generations where the accelerated accumulation (initiation of death) started (blue). (F) Probability of death of cells born with different amounts of aggregate. Gray arrowheads indicate data points whose deviations from the population death probability (*p* = 1.15 × 10^-2^) are statistically significant (binomial test at the significance level α = 0.05. See Materials and Methods for details). Data points within gray-shaded area do not satisfy the commonly employed rule of *Np* ≥ 5 for appropriate normal approximation (*N*, the number of samples), and are excluded from the tests. Error bars show standard errors. The same rule applies to the arrowheads, gray-shaded area, and error bars in panels (G), (I), and (J) below. (G) Probability of initiating cell death (= observing a kink) for cells born with different amounts of aggregate. (H) Distributions of aggregation age for all of the ROIs (gray), at death points (red), and at kinks (blue). (I) Death probability at each aggregation age. (J) Probability of observing a kink at each aggregation age.

Due to the occurrence of accelerated accumulation in many extinct lineages before deaths, the distribution of aggregates at the death points was shifted toward greater values (Fig. 6D). However, the distribution of the amount at the kink points was very close to that for the total population (Fig. 6D), which suggests that large aggregate amounts are not required for initiating the process of dying. To reinforce this observation, we counted the numbers of cells that exhibited deaths and kinks for given ranges of aggregate amount (Fig. 6E), and evaluated the probability of death (Fig. 6F) and of starting accelerated accumulation (Fig. 6G). The results showed that the probability of commencing accelerated accumulation did not increase with the aggregate amount, although that of observation of cell deaths was elevated, which again suggests that the aggregate amount is not causative of initiation of the dying processes.

We also investigated whether retention of protein aggregates increased the probabilities of death and of starting accelerated accumulation by counting the numbers of cells that showed deaths and kinks for given aggregation age (Fig. 6H). We found that both probabilities were nearly constant, irrespective of aggregation age (Fig. 6I and 6J). The death probabilities for aggregation age < 3 generations were slightly lower than the total death probability (1.15 × 10^-2^ per generation) (Fig. 6I) possibly due to an identified lag of few generations before death, after the onset of accelerated accumulation (Fig. 6C). Indeed, the probabilities of commencing accelerated accumulation were equally high for these small aggregation-age generations (Fig. 6J). Overall, our data suggest that Hsp104-associated protein aggregation is unlikely to play a major role in initiating the dying process. Our results, however, do not exclude the possibility that rapid accumulation of protein aggregate might accelerate completion of the dying processes post-onset.

### Protein aggregation induced by oxidative stress does not result in replicative aging

It has been suggested that fission yeast ages upon stress treatment, and inheritance of large protein aggregate results in increased death probability [11]. To see if these aging phenotypes are also observed in our system, we transiently treated cells with hydrogen peroxide (H_2_O_2_), a commonly-used oxidative stressor, and monitored cell division/death kinetics along with protein aggregation dynamics. As expected, cells immediately ceased to divide upon stress treatment (Fig. 7A, around *t* = 6,000 min). After removal of the stress, there was a lag (around *t* = 6,000–6,500 min) before cells resumed dividing. Strikingly, once cells started to grow again, the division rate was almost the same as that previously observed under unstressed conditions (Fig. 7A). The survival curve in Fig. 7B revealed an increase in death rate upon stress treatment (∼10% cells died during an hour of exposure to hydrogen peroxide). Although the recovery was slower than division rate, the death rate also returned to the normal level seen in the unstressed condition. We did not observe the progressive increase of generation time after stress removal, one of the hallmarks of replicative aging; marked increase of generation time was seen only in the first generation after stress removal (Fig. 7C). These results suggest that the apparent deterioration in cellular growth/death is a transient response to the stress, and not a manifestation of aging.

**Fig 7.**
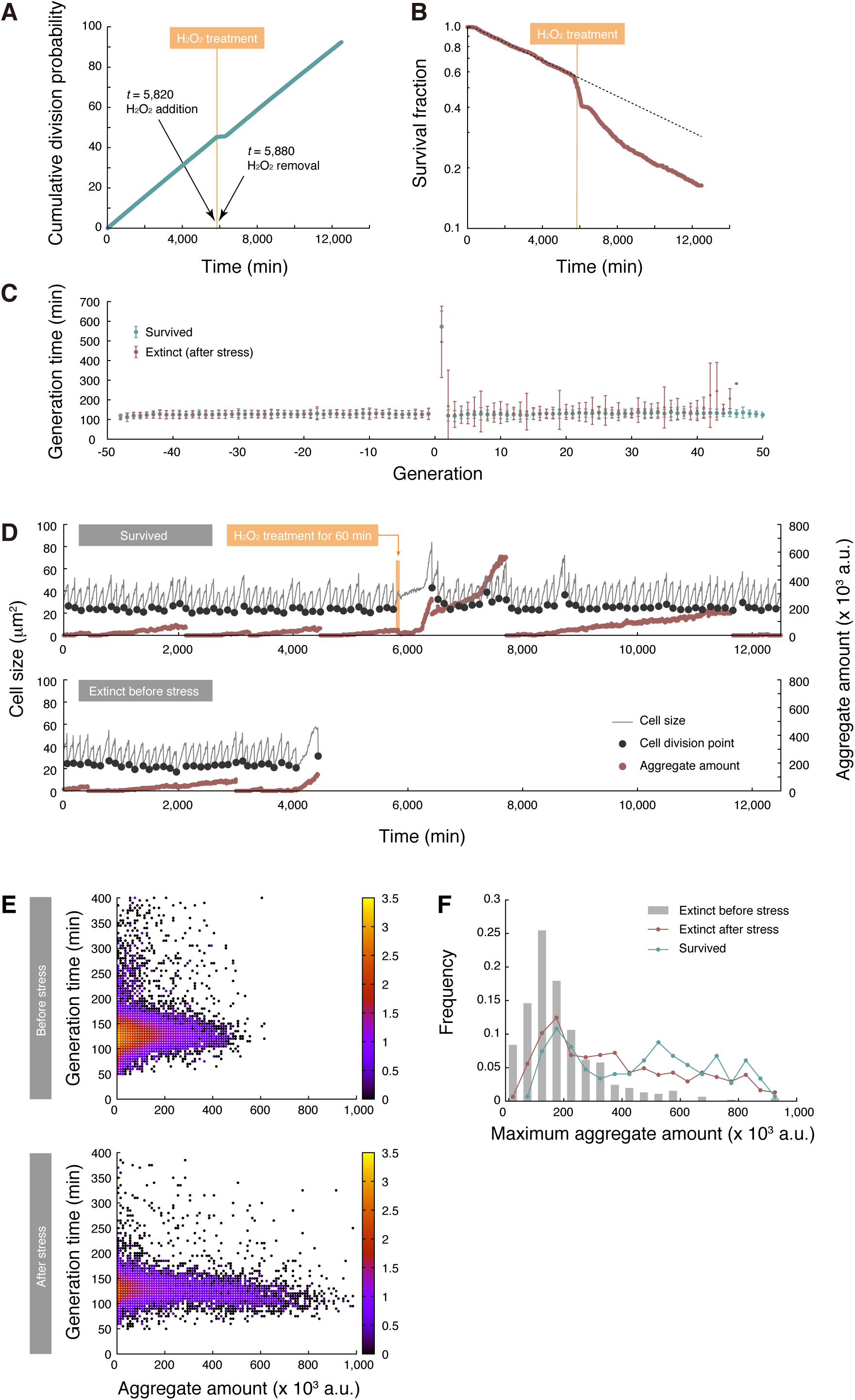
Oxidative stress does not induce replicative aging. (A) Cumulative division probability. Mean generation time, estimated as a reciprocal of the slope of the line, was 130 min before stress, and 134 min after recovery, respectively. Orange thick line represents the period of H_2_O_2_ treatment (60 min). (B) Survival curve before and after stress treatment. The broken line represents a predicted survival curve without stress treatment. (C) Transitions of mean generation time before and after stress. Positive and negative generations indicate the generations after and before H_2_O_2_ treatment, respectively. Lineages that expired before stress treatment were excluded from the analysis. (D) A representative example of a lineage that survived until the end of measurement (top), and a lineage that died before stress treatment (bottom). Stress treatment induces much higher levels of aggregation than in normal conditions. (E) Density maps for the relationship between generation time and aggregate amount. (Top) Before stress treatment. (Bottom) After stress treatment. Density of points is represented in color (log_10_[Counts]). (F) Distributions of maximum aggregate amount on a lineage. Gray color indicates the lineages that were extinct before stress; red for the lineages that were extinct after stress; blue for the survived lineages.

We next asked how protein aggregation dynamics are related to the stress response. As reported earlier, we observed that oxidative stress enhanced protein aggregation, and many lineages accumulated aggregate to high levels not attainable in normal conditions (Fig. 7D). When the cells re-entered division cycles after stress removal, the large amount of aggregate persisted (and even continued to grow in some cases). Strikingly, even such significant amounts of aggregate did not affect generation time (Fig. 7E). In non-stressed conditions, the amount of aggregate did not exceed 400 (× 10^3^ a.u.) in 90% of extinct lineages, whereas approximately 40% of the lineages that survived the stress treatment until the end of measurement experienced more aggregation than 400 (× 10^3^ a.u.) (Fig. 7F). These results further support that the absolute amount of protein aggregate does not determine growth kinetics, nor cell fates.

### Ectopically induced protein aggregation does not affect cellular growth or death

To further examine if protein aggregation can result in cellular aging and/or cell death in *S. pombe*, we ectopically expressed a truncated version of the orthoreovirus aggregation-prone protein, μNS, which was N-terminally tagged with mCherry or mNeonGreen for visualization by fluorescent microscopy [32–34]. We confirmed that μNS formed aggregates in the cytoplasm of *S. pombe*, which are detectable as bright foci in many cells (Fig. 8A). mCherry-μNS and Hsp104-GFP foci did not co-localize, which indicates that not all protein aggregates were associated with Hsp104 (Fig. 8A and Movie S5). Formation and segregation dynamics of mNeonGreen-μNS aggregate were similar to those of Hsp104-GFP, other than that accelerated accumulation before cell death was not observed (Fig. 8B, S8A, and Movie S5). Generation time was not correlated with either aggregate amount or aggregation age (Fig. 8C), and distribution of μNS aggregate amount at death points was almost identical to that at the termination of the measurements for the survived lineages (Fig. 8D). These results suggest that, as noted for endogenous Hsp104-associated protein aggregation, induction of ectopic protein aggregation causes no functional loads on *S. pombe* cells in terms of the rates of cell division and initiation of death.

**Fig 8.**
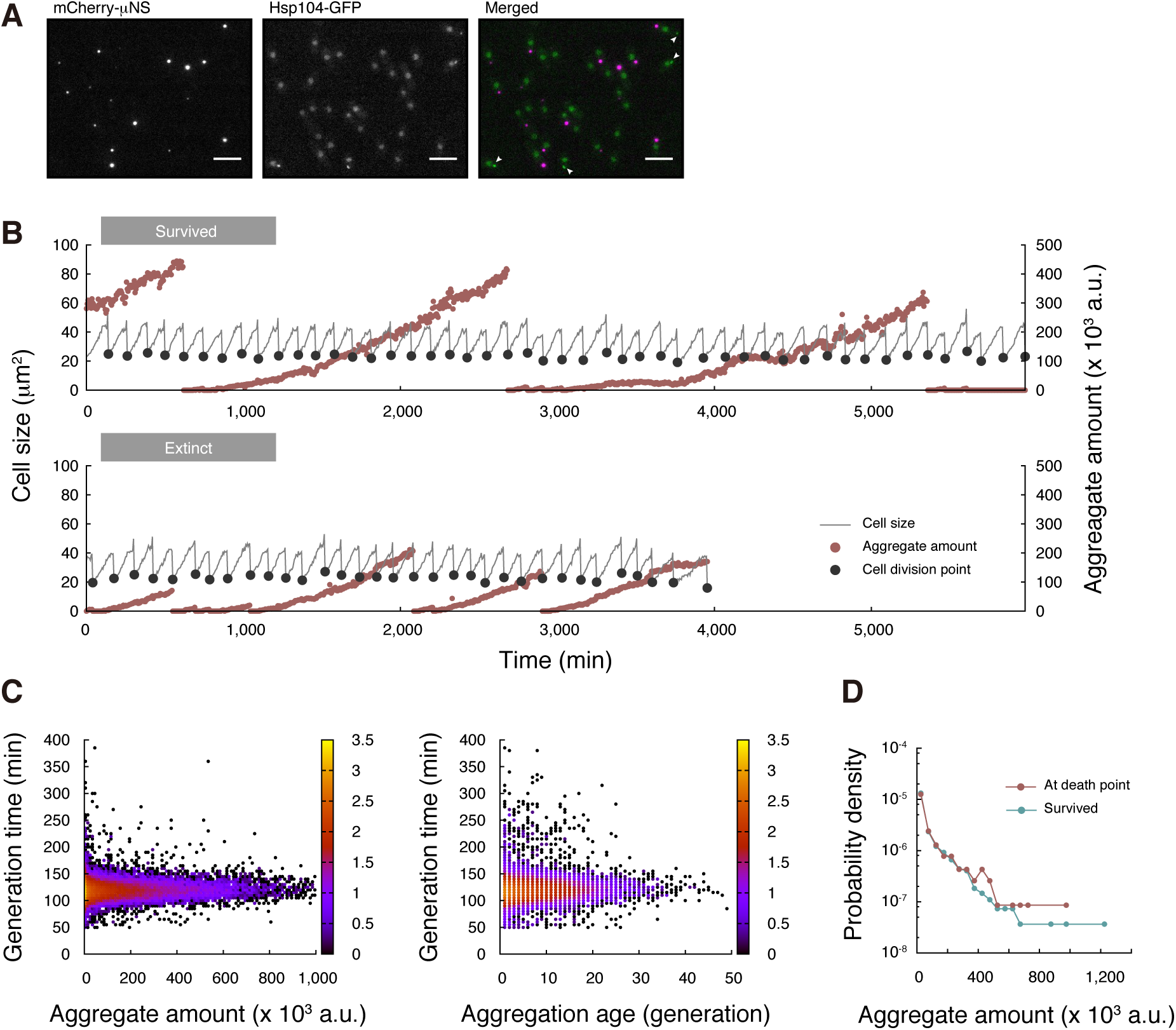
Ectopic expression of aggregation-prone protein does not affect cellular growth and death. (A) μNS and Hsp104 form distinct types of aggregate. mCherry-μNS under the control of the *nmt1P41* promoter was co-expressed with Hsp104-GFP in EMM lacking thiamine. (Left) Cytoplasmic foci of mCherry-μNS aggregates. (Middle) Hsp104-GFP aggregates are observed as cytoplasmic foci. In most cases, nuclei are also diffusely stained. (Right) Merged image. No co-localization of μNS and Hsp104 is observed. White arrowheads indicate cytoplasmic Hsp104-GFP foci. Scale bars indicate 10 μm. (B) Representative dynamics of μNS aggregate amounts shown with cell division cycles. (C) No correlation between generation time and aggregate amount (left), and aggregation age (right). (D) Distributions of aggregate amounts at death points (red) and the end point of the survived lineages (blue).

### Deletion of ***hsp104^+^*** gene does not result in aging

Finally, we examined if *hsp104*^+^ gene disruption results in replicative aging in the old-pole cell lineages. As expected, *hsp104*Δ strains were sensitive to heat shock (Fig. 9A). The deletion, however, did not affect the status of μNS aggregate in terms of both inheritance duration and distribution of the amount (Fig. S8A and S8B). Likewise, generation time remained non-correlated to aggregation (Fig. S8C), and distribution of aggregate amount at death points was similar to that at the end points of the survived lineages (Fig. S8D). Division- and death rates for the deletion mutant were essentially identical to those in the wild type strain (Fig. 9B and 9C). Taken together, our data suggest that regulation of protein aggregation by Hsp104 is not critical for avoiding replicative aging in the old-pole lineages of *S. pombe*.

**Fig 9.**
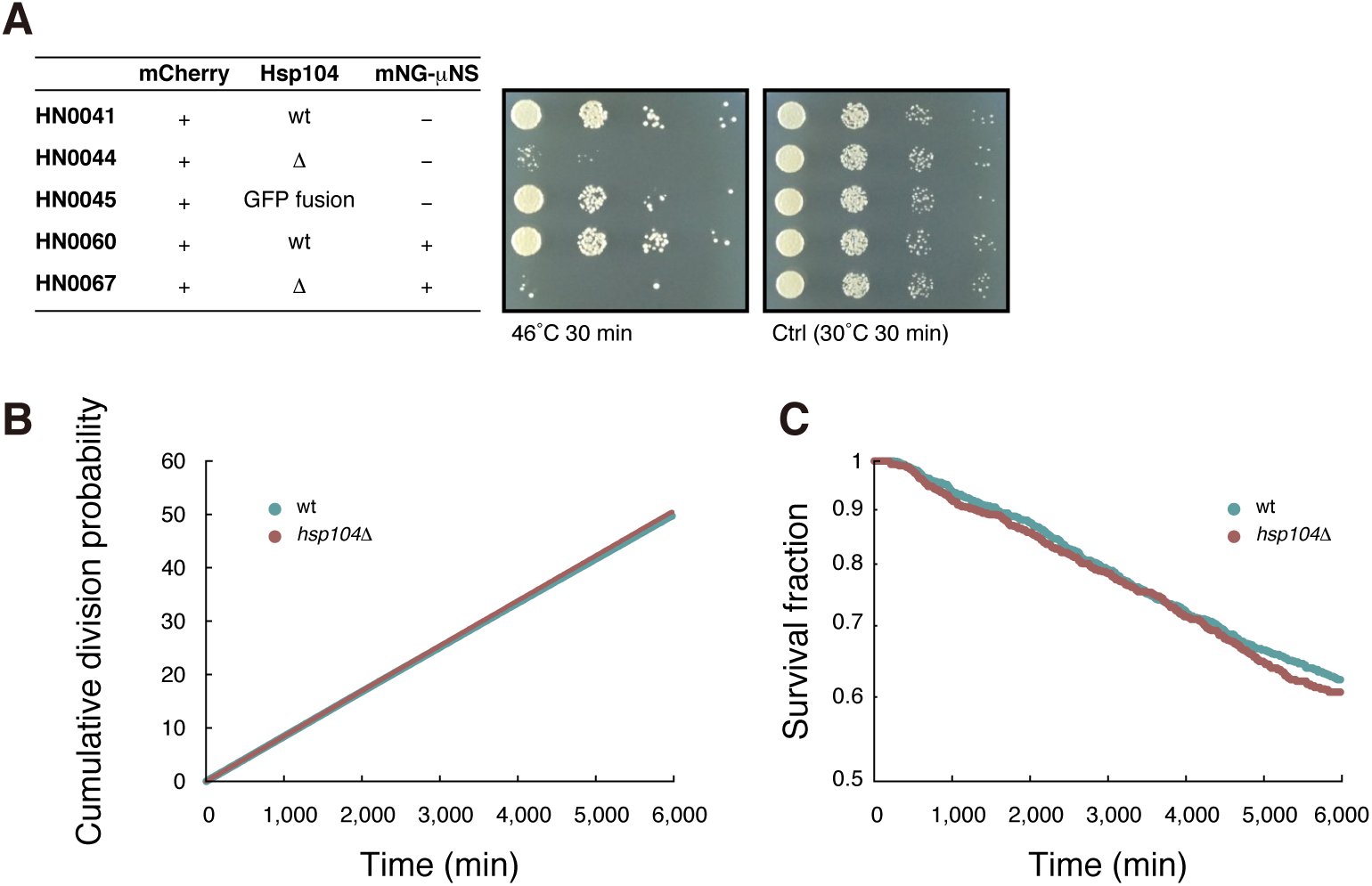
Disruption of *hsp104*^+^ does not cause replicative aging. (A) Spot assays showing that *hsp104Δ* is sensitive to heat shock. (B) The cumulative division probability plotted against time. Division rates estimated by the slope of the fitted lines are (8.319 ± 0.001) × 10^-3^ min^-1^ for wild type and (8.448 ± 0.001) × 10^-3^ min^-1^ for *hsp104Δ*, respectively. (C) Survival curves. Death rates estimated by exponential fitting are (8.55 ± 0.01) × 10^-5^ min^-1^ for wild type and (8.85 ± 0.01) × 10^-5^ min^-1^ for *hsp104Δ*, respectively.

## Discussion

In this study, we investigated the growth and death of *S. pombe* under various favorable culture conditions, and demonstrated the lack of replicative aging in old-pole cell lineages. While being free from replicative aging, the cells died abruptly, and our extensive quantification revealed trade-offs between reproduction rate and survival probability. Furthermore, we showed that Hsp104-associated protein aggregation, often regarded as an aging factor, did not have a significant effect on growth and initiation of the dying processes, not only under normal conditions, but also when stressed, where a large quantity of protein aggregate was induced.

### Replicative aging in fission yeast

The observation that both division and death rates were constant over tens of generations strongly suggested that the fission yeast old-pole cell lineages were free from replicative aging (Fig. 1 and Fig. 2). This endorses the conclusion of a recent report on the absence of replicative aging in fission yeast in favorable conditions [11]. However, we cannot formally exclude the possibility that aging of fission yeast occurs over a longer time-scale than the observed durations in our experiments and in other recent work [11,27]. Regarding the discrepancy with the earlier reports, where replicative aging was suggested [8,10], it is of note that in the cited mortality assays, the populations became extinct within 20 generations while about 80% of cells survived at that generation in our experiments (Fig 2E). This implies that the physiological states of the observed cells in the experiments reported elsewhere were significantly different from those in ours.

Our experiments involving oxidative stress exposure in the microfluidic device revealed that generation time and death rate reverted to normal after removal of the stressor, although most old-pole cells continued to inherit and accumulate even more Hsp104-associated protein aggregates (Fig 7). This result is inconsistent with the previous report [11] and the widely accepted notion that *S. pombe* ages under stressful conditions. The lack of progressive increases in generation time suggests that the observed increases of generation time, death rate, and the amounts of Hsp104-associated protein aggregation are responses provoked by the stress exposure, not the signature of aging. Therefore, aging under stressful conditions might not be a general trait in *S. pombe*.

The lack of noticeable aging in *S. pombe* old-pole lineages contrasts with that in *E. coli* cultured under favorable growth conditions in the Mother Machine, in which division rates of old-pole cells were stable, but death rates increased over generations [7]. It is interesting to note that the modes of cell wall syntheses at poles are quite different between the two symmetrically dividing microorganisms. *S. pombe* employs polar growth and newly synthesized cell wall materials are exclusively incorporated at the ends of the cells [35–37], whereas in *E. coli*, the cylindrical part of the cell grows and cell walls at poles are thought to be metabolically inert and unable to avoid deterioration [38–40]. Thus, for fission yeast we do not have an *a priori* reason to believe that old-pole lineages should undergo senescence. Rather, new-pole lineages that inherit a larger proportion of old lateral cell walls and a birth scar (due to the delay for the new-pole end to initiate growth after division [41]) might be subject to aging. It should be stressed that our results do not rule out the existence of any forms of lineage-specific aging in fission yeast, e.g. lineages that inherit damaged (carbonylated) proteins and/or birth scars [10]. Since the Mother Machine allows tracking of only old-pole cell lineages, new methods are required to track such cell lineages within proliferating populations.

Our results might also imply that symmetrically dividing unicellular organisms could escape aging if they possess efficient damage repair mechanisms. On the other hand, systematic asymmetric partitioning of most cellular components should occur in non-symmetrically dividing unicellular organisms, resulting in consistent unequal partitioning of the damaged materials to specific lineages, associated with cellular morphological features such as cell size. This might underlie the empirical fact that aging is more prominent in asymmetrically dividing unicellular organisms.

### Protein aggregation and cellular growth/death

A common perception is that protein aggregate accumulates during the aging process or the stress response, and cells die catastrophically when the aggregation load exceeds the cellular capacity. Our data, however, indicated that Hsp104-associated protein aggregate is also formed in aging-free cell lineages (Fig. 4 and Movie S4). We showed that neither the aggregate amount nor the retention time affected the generation time (Fig. 5). In addition, we demonstrated that cells transiently exposed to oxidative stress could promptly resume normal growth, even in the presence of unusually large amounts of protein aggregate induced by such stress (Fig. 7). The commonly-observed correlation between protein aggregation and cell death is most likely explained by the accelerated accumulation of aggregate a few generations before cell death. The commencement points of accelerated accumulation appear to specify the initiation points of the dying processes because the other abnormalities of mCherry expression levels and cellular morphology started around the same time. The initiation of the dying process occurred irrespectively of aggregate quantity, which argues against the concept that there is an absolute threshold in protein aggregation burden for triggering cell death. The results also suggest that retention of the aggregates did not elevate cell death probability (Fig. 6H–J). Overall, our data indicate that Hsp104 foci do not reflect gradual deterioration of proteostasis in the cells, and thus cannot be used as the sole molecular marker for cellular senescence in fission yeast.

Together with the results regarding Hsp104-associated protein aggregation, our experimental results involving the ectopic aggregation-prone protein, μΝS, suggest that *S. pombe* is, in fact, highly tolerant of protein aggregation loads (Fig 8), and that their adverse effects on growth and death and their relation to aging may have been significantly overestimated.

It should be noted that there can be multiple types of protein aggregation with different constituents and formation/degradation dynamics, such as stress foci/Q-bodies/CytoQ, IPOD, JUNQ/INQ, and age-associated deposits, as reported for budding yeast [17,42–44]. Hsp104 is enriched in stress foci, IPOD, and age-associated deposits, but typically not in JUNQ/INQ [17]. Although Hsp104 is assumed to represent total protein aggregates in fission yeast [18], detailed classification and characterization of protein aggregates in this organism are still lacking. Therefore, future studies should clarify to what extent Hsp104-associated protein aggregates capture the behaviors of total protein aggregates in cells.

### Trade-off between reproduction and survival

We found that as the cell division rate elevated, the death rate increased in a linear fashion (Fig. 3A). Although a simple extrapolation of the linear trend predicts immortality of single cells when the division rate is below *r*^min^, we have not been able to experimentally achieve stable growth with such a low division rate under our current measurement setup. The slow growth of cells with a division rate close to, or even smaller than, *r*^min^ could be achieved by applying stressors, such as high/low temperatures, nutrient limitation, drug exposure, or harsh chemical conditions (e.g., extreme redox environments, high/low osmolarity). However, stress responses would render internal cellular states different from those in non-stressed conditions. We speculate that this linear trend is a hallmark of a balanced growth state, rather than a universal constraint on cell division and death rates in any environment.

Why might death rate be positively-correlated with division rate? In line with the historical free radical theory of aging, it could be contemplated that faster growth with higher metabolic rates generates greater oxidative stress and/or toxic metabolic wastes that result in cell death [45]. Although a substantial body of work supports the notion that reactive oxygen species (ROS) underlie aging and/or the reproduction-survival trade-off, skeptical views have also been proposed in recent studies [46–48]. In addition, caution should be exercised when applying these theories to aging and/or life history, given the lack of aging before death in our observations. Another, and not mutually exclusive, possibility is that cells might allocate energy and resources to growth and division at the expense of maintenance mechanisms such as DNA repair, protein quality control, and stress responses. Indeed, the expression of stress-induced genes is negatively correlated with growth rate in budding yeast [49], and enhanced stress resistance in slower growing cells has been reported [50,51]. Demetrius proposed that it is neither metabolic rate, nor specific metabolites, but rather the stability of the entire metabolic system that determines lifespan [52]. However, how external environments (i.e. temperature and/or nutrition status) affect the robustness of steady-state levels of metabolites to stochastic fluctuations in metabolic processes is still poorly understood and would be an interesting topic for future research.

### Causes of cell death

In general, death can be triggered by both external and internal cues [53]. Because the microfluidic system ensures stable culture conditions by continuous supply of fresh medium, cellular deaths observed in this study are most likely caused by internal signals, although we cannot exclude the possibility that subtle fluctuations of local environments around cells had some impacts on death. DNA lesions are unlikely to be a major cause of death because the extensive elongation in cell length, a characteristic phenotype of DNA damage response [54], was observed in only ∼20% cases (Fig. S4E). ROS are involved in many examples of programmed cell death from yeasts to humans [55–57], and thus are at least one of the significant candidates to be examined. Accordingly, it would be of great interest to visualize the dynamics of mitochondria and/or peroxisomes in our experimental system. A very recent study has reported that Sir2p (Silent Information Regulator 2: a highly-conserved NAD^+^-dependent deacetylase) overexpression or inhibition of the TOR pathway (Target Of Rapamycin: a highly-conserved phosphatidylinositol kinase-related protein kinase that is responsible for nutrient responsive growth regulation) by rapamycin decreases death rates in *S. pombe* [27]. Involvement of a variety of biological processes implicated by such observations, including heterochromatin regulation [58,59], asymmetric partition of damaged proteins [10], and nitrogen-starvation responses [60–62], should be investigated in future research.

At the present time, however, we do not understand if sporadic death is dependent on any specific molecular mechanism. For example, intrinsic fluctuations in gene expression and/or biochemical reaction rates may well differentially result in catastrophic alterations to cellular physiology. A variety of fluorescent biosensors for more global physiological states, such as intracellular pH, ATP, NAD^+^/NADH, molecular crowding, and second messengers—cAMP, DAG, Ca^++^ etc. would be valuable and informative to infer the causes of cell death [63–68]. We hope that our simple but powerful microfluidics approach can contribute to the detailed study of the mechanisms of cell mortality, and a deeper understanding of limitations in cellular homeostasis, one of the significant questions in biology.

## Materials and Methods

### Microfluidic device fabrication

The microfluidic device was fabricated by standard photolithography techniques [69–71] (Fig. S1B). The CAD designs, created using ZunoRAPID software (Photron), were printed on photoresist-coated chrome-on-glass masks (CBL4006Du-AZP, Clean Surface Technology) using a laser-drawing system (DDB-201-TW, Neoark). The UV-exposed regions of the photoresist (AZP1350) were removed by NMD-3 (Tokyo Ohka Kogyo) and the exposed chromium was etched by MPM-E350 (DNP Fine Chemicals). After removing the remaining photoresist layer using acetone (Wako), the masks were rinsed with MilliQ water and air-dried.

An SU-8 mold for the PDMS (polydimethylsiloxane) device was made on a silicon wafer (ϕ= 76 mm, P<100>, resistance 1 to 10 W cm, thickness 380 μm; Furuuchi Chemical) in two steps. First, to make observation channels, the wafer was coated with SU-8 3005 (Nippon Kayaku) using a spin-coater (MS-A150, Mikasa) at 500 rpm for 10 sec, then 4,000 rpm for 30 sec. After soft baking for 2 min at 95°C, the photoresist was exposed to UV using the mercury lamp of a mask-aligner (MA-20, Mikasa) at 22.4 mW/cm^2^ for 12 sec. Post-exposure baking was performed at 95°C for 3 min, followed by exposure to SU-8 developer (Nippon Kayaku) and a 2-propanol (Wako) rinse. The same procedure was repeated to fabricate trenches using SU-8 3025 (Nippon Kayaku) at 500 rpm for 10 sec, then 2,000 rpm for 30 sec. Soft baking was performed at 95°C for 10 min, followed by UV exposure at 22.4 mW/cm^2^ for 16 sec and post-exposure baking at 95°C for 10 min.

The PDMS base and curing agent (Sylgard 184) were mixed at a ratio of 10:1, poured onto the SU-8 mold in a container, and de-gassed using a vacuum desiccator. Curing was performed at 65°C overnight. The device was peeled from the mold, washed briefly in ethanol with sonication, and then air-dried. After punching two holes in the device to connect the inlet and outlet tubes, the surfaces of the device and a coverslip (24 × 60 mm, thickness 0.12–0.17 mm, Matsumani) were activated using a plasma cleaner (PDC-32G, Harrick Plasma) and bonded together. Finally, the inlet and outlet tubes were inserted into the holes (see also Fig. S1).

### Fission yeast strains

HN0025, which expresses mVenus under the control of a constitutive *adh1* promoter (*h^-^ leu1-32*::*leu1^+^-Padh1-mVenus*), was constructed by transforming HN0003 (*h^-^ leu1-32*) with the *Not* I-digested fragment of pDUAL-mVenus, generated by replacing the *Sph* I*-Cla* I fragment of pDUAL 13G10 with the *Padh1-mVenus-nmt1 terminator* cassette. HN0041 was constructed in the same manner as HN0025 except that that mVenus fragment was replaced with mCherry, and used as a parental strain to establish HN0045. To generate HN0045, the GFP tag was fused at the C-terminus of Hsp104 by PCR-based gene targeting [72]: the 3’ end of the ORF and 3’ UTR region of *hsp104*^+^ were assembled with the GFP(S65T)-*kan^R^* fragment on the pFA6a-GFP(S65T)-KanMX6 to generate a targeting module. For the strain depicted in Fig. S1D (HN0034), the *Padh1-mCherry* cassette was integrated at the *ade6*^+^ locus. To generate HN0060, *Ptef-mNeonGreen-μNS* cassette was first cloned into pDUAL vector, and then a *Not* I-digested fragment was integrated to the *leu1-32* locus. Note that the μNS is a truncated version (1411–2163) that corresponds to amino acids 471–721. Genomic DNA of bacterial strain CJW4617 [34], (a kind gift from Dr. M. Nibert at Harvard Medical School and Dr. C. J. Wagner at Yale University), was used as a template for PCR-cloning of the μNS cDNA. To disrupt *hsp104*^+^, the host strains were transformed by a PCR-amplified deletion cassette, and G418-resistant clones were selected. A complete list of the strains and PCR primers used in this study can be found in Table S4. The pDUAL 13G10 strain was provided by Riken BRC, a member of the National Bio-Resources Project of the MEXT, Japan [73,74].

### Confocal microscopy

Confocal fluorescence microscopic images of yeast cells in the microfluidic device (Fig. S1D) were acquired using a Nikon Ti-E microscope equipped with a laser scanning confocal system, equipped with a 60 × objective lens (Plan Apo λ N.A. 1.4, Nikon) and under oil immersion. The resolution along the Z-axis was 0.15 μm, and 335 images (corresponding to approximately 50 μm in height) were taken. 3-dimensional reconstructions of the images were achieved using an ImageJ 3D viewer plug-in.

### Long-term time-lapse experiments

For long-term time-lapse measurements, 10 mL of a log-phase culture of yeast cells at 28–34°C in YE containing 3% glucose or EMM containing 2% glucose was concentrated 50 fold by centrifugation and injected into the microfluidic device using a 1 mL syringe (Terumo). Cells were loaded into the observation channels by gravity, simply slanting the device. The loading procedure typically took a couple of hours, during which time the cells often entered into an early stationary phase in response to the highly crowded environment, resulting in a time lag before stable growth was achieved. The device was supplied with appropriate medium supplemented with a low concentration (10 μg/mL) of ampicillin sodium (Wako) to minimize the risk of bacterial contamination. Note that ampicillin has no effect on fission yeast growth. The flow rate was 10–15 mL/h. For transient oxidative stress treatment, the medium was changed to YE containing 2 mM hydrogen peroxide for 1 hour, then switched back to YE.

We used a Nikon Ti-E microscope with a thermostat chamber (TIZHB, Tokai Hit), 40 × objective (Plan Apo λ N.A. 0.95, Nikon), cooled CCD camera (ORCA-R2, Hamamatsu Photonics), and an LED excitation light source (DC2100, Thorlabs). Several cell divisions were allowed before initiating measurements. Micromanager software (https://micro-manager.org/) was used for fluorescence and/or bright field image acquisition. The time-lapse interval was 3 min (for the experiments described in Fig. 1–3), or 5 min (for the experiments described in Fig. 4–9). Exposure times were 400 ms (for mVenus), 200 ms (for GFP), 100 ms (for mCherry), and 10 ms (for bright field).

### Single-cell lineage tracking and division/death points identification

The acquired fluorescence images were converted into binary images using a custom-written OpenCV program. The binary images were used to identify cellular regions or ROIs, and lineage tracking (relating ROIs along lineages) was performed using a customized ImageJ macro. Transition between ROIs along each lineage was analyzed to mark cell division points where an ROI area suddenly decreased more than 1.5 fold. To mark cell death points, two criteria were employed: 1) if there was no division during a 360-min window, then the beginning of the window was defined as a death point, and 2) if there was a profound (more than 1.75 fold) decrease in fluorescence during a 30-min time window, then the beginning of the window was defined as a death point. We confirmed that the decay curve of surviving cell lineages obtained using these death criteria quantitatively concurred with that obtained by manual image inspection (Fig. S4A). In the data set used in Fig. 6–9, we examined all of the cell-size trajectories by eye and manually marked death points so as to ensure confidence in the data.

The death onset points (= kinks on the aggregate amount trajectories) were identified by manually inspecting the aggregate amount trajectory plots for all of the 541 extinct lineages. Aggregate amount/age at the kinks and generations to die after the onset of the dying process were subsequently recorded using a custom-developed ImageJ-plugin.

### Estimation of the population doubling time from the distribution of intrinsic generation time

Generation time distributions obtained by following old-pole cell lineages in the Mother Machine represent intrinsic cellular division properties when two sister cells are physiologically indistinguishable. In standard batch cultures, however, selection occurs because of heterogeneities in cellular generation times, which cause the population doubling time (*T_d_*) to be smaller than the mean generation time 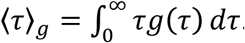. Here *g*(*τ*) is the probability density function of an intrinsic generation time distribution in a given culture condition. When cells randomly and independently determine their generation times according to *g*(*τ*), the population growth rate, Λ = ln2/*T_d_*, must satisfy the Euler-Lotka equation [28–30],

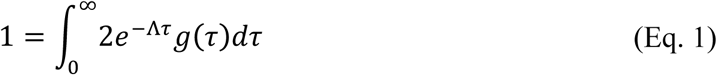

Equivalently,(Eq.1) can be rewritten as

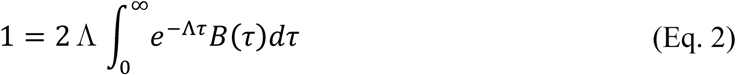
Where 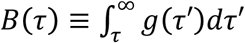 is the survival function (complementary cumulative distribution) of *g*(*τ*), which represents the probability of a newly divided cell remaining undivided until age τ. One can show that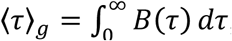, and then 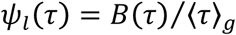 becomes a probability density function, and can be interpreted as “age distribution” along cell lineages. Therefore, Eq. 2 can be also expressed as

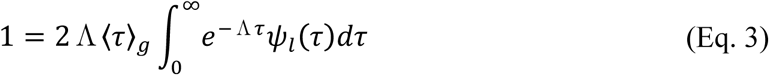

We numerically estimate Λ based on a discretized version of Eq. 3, i.e.,

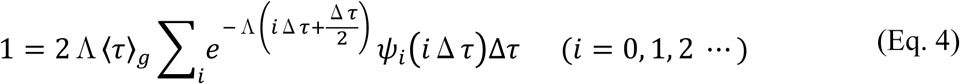
where Δ*τ* is the time-lapse interval. 〈*τ*〉*_g_* was calculated as 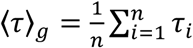, where *τ_i_* is the generation time and *n* is the number of samples, and the probability distribution of age is calculated as 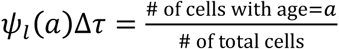.

### Division rate, death rate, and expected life span of fission yeast

We calculated division rate *r* as

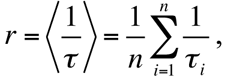
where *τ_i_* is the generation time and *n* is the number of samples.

We estimated death rate *k* from the decay curve by the least squares fitting (*t* versus ln(*N*(*t*)), where *t* is time and *N*(*t*) is the number of surviving lineages at *t*). The expected value of “time to death” is thus 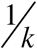. We calculated “expected life span” in units of generation as 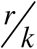.

### Error estimation of division and death rates

The standard errors for the division rates were calculated as

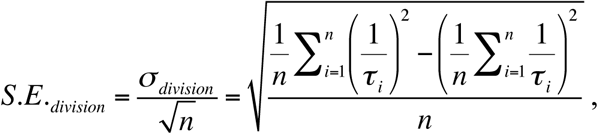
where *τ_i_* is the generation time and *n* is the number of samples. ±2S.*E._division_*ranges were shown as error bars in Fig. 3A and 3B.

To evaluate errors in the death rate estimations, we produced simulated decay curves of the surviving fraction using parameters (death rate, initial cell number, and observation period) specific to each experiment. The simulation was repeated 5,000 times for each environment, and the death rate was obtained from a simulated survival curve in each run. The standard deviation of the determined death rates was calculated as *σ_death_*, and the ±2*σ_death_* ranges were shown as error bars in Fig. 3A. Errors in expected life span (σ*_lifespan_*) were calculated using the error propagation rule.

### Statistical evaluation of death (or death onset) probability

We first estimated death probability per generation *p_0_* to be 1.15 × 10^-2^ from the survival curve. In Fig. 6F and I, death probability *p* for each aggregate amount or aggregation age was then tested using a binomial test for the two-tailed null hypothesis *H_0_* : *p* = *p_0_* at the significance level = 0.05.

For the onset probability of accelerated accumulation, we set the null hypothesis to be *H_0_* : *q* = 0.79 *p_0_* = 9.09 × 10^-3^ based on our observation that a clear kink in the protein aggregation dynamics was detected in 79% of the extinct lineages, and implemented binomial testing at the significance level = 0.05 (Fig. 6G and 6J).

## Author Contributions

Conceptualization, H.N. and Y.W.; Investigation, H.N.; Writing – Original Draft, H.N. and Y.W.; Writing – Review & Editing, H.N. and Y.W.; Funding Acquisition, H.N. and Y.W.

## Acknowledgments

We thank Kunihiro Ohta and Takatomi Yamada for the kind gift of fission yeast strains, Max Nibert and Christine Jacobs-Wagner for the kind gift of bacterial strains, Riken BRC for plasmids, Reiko Okura for technical assistance, and members of the Wakamoto lab for in-depth discussions.

